# Naturally Produced Lovastatin Modifies the Histology and Proteome Profile of Goat Skeletal Muscle

**DOI:** 10.1101/581439

**Authors:** Teik Kee Leo, Sani Garba, Danmaigoro Abubakar, Awis Qurni Sazili, Su Chui Len Candyrine, Mohammad Faseleh Jahromi, Yong Meng Goh, Ron Ronimus, Stefan Muetzel, Juan Boo Liang

**Author notes:** Corresponding author (JBL).

## Abstract

Enteric methane formation in ruminants is one of the major contributors to climate change. We have reported that supplementation of naturally produced lovastatin reduced methane emissions in goats without adversely affecting rumen fermentation and animal performance, except that at higher level, lovastatin can have a negative effect on the palatability of the formulated diet. As statins are associated with the development of muscle-related adverse effects at higher than recommended therapeutic doses, this study was conducted to examine the effects of lovastatin on the histology and proteome profile of goat skeletal muscle. A total of 20 intact male Saanen goats were randomly assigned in equal numbers to 4 groups, and fed with a total mixed ration containing 50% rice straw, 22.8% concentrates and 27.2% of various proportions of untreated or treated palm kernel cake (PKC) to achieve the target daily intake levels of 0 (Control), 2 (Low), 4 (Medium) or 6 (High) mg lovastatin/kg body weight (BW). Histological examination discovered that the longissimus thoracis et lumborum muscle of animals from the Medium and High treatment groups showed abnormalities in terms of necrosis, degeneration, interstitial space and vacuolation. Western blot analysis conducted on the myosin heavy chain showed that the immunoreactivity of myosin heavy chain in the High treatment group was significantly lower than the Control, Low and Medium treatment groups. Comparisons between control and lovastatin-treated groups demonstrated that lovastatin supplementation induced complex modifications to the protein expression patterns of the longissimus thoracis et lumborum muscle of the goat. There were 30, 26 and 24 proteins differentially expressed in Low, Medium and High treatment groups respectively, when compared to the Control group. Supplementation of lovastatin down-regulated proteins involved in carbohydrate and creatine metabolism, indicative of reduced energy production, and may have contributed to the skeletal muscle damage. Supplementation of naturally produced lovastatin induced muscle damage in longissimus thoracis et lumborum muscle of goats with increasing dosages, particularly at 6mg/kg BW. In addition, proteomics analysis revealed that lovastatin supplementation induced complex modifications to the protein expressions of skeletal muscle of goats which may have contributed to the observed skeletal muscle damage. Present study suggested that supplementation of naturally-produced lovastatin at 6mg/kg BW could adversely affecting health and wellbeing of the animals.

## Introduction

Methane gas is one of the major greenhouse gases that is contributing to climate change. Livestock production has been reported to contribute approximately 18% of methane emissions and 9% of carbon dioxide production [1], which results primarily from the enteric fermentation of feeds [2]. Enteric methane formation results from the activity of complex interactions of anaerobic bacteria which together enable degradation of ruminant feeds and methanogenic archaea which help remove metabolic hydrogen in the rumen [3]. Despite the importance of methanogenesis in maintaining low partial pressure of hydrogen required for efficient ruminal fermentation, the formation of methane also represents 2-12% loss of gross dietary energy [4]. Hence, extensive research efforts are focused on the development of strategies to modify ruminal fermentation for reduction of methane emissions [5] as well as better feed utilization [6].

Among the potential strategies for mitigating methane emissions, supplementation of feed additives such as ionophores [7], organic acids [8], fatty acids [9], methyl coenzyme M reductase inhibitors [10], vaccine [11] and oils [12] have been extensively researched. However, many of these strategies have limited application due to their negative effect(s) on human and animal health, animal performance parameters and economical acceptance [13]. Supplementation of naturally produced lovastatin is a promising approach for mitigating methane emissions.

Lovastatin (C24H36O5, M.W. 404.55) is a secondary product during the secondary phase (idiophase) of fungal growth. It is a competitive inhibitor of 3-hydroxy-3-methylglutaryl coenzyme A (HMG-CoA) reductase, which is a key enzyme in the cholesterol production pathway [14]. Inhibition of HMG-CoA reductase will mediate the suppression of cholesterol synthesis and cell membrane formation in methanogenic Archaea. A previous study has shown that significant reduction in growth and activity of methanogenic Archaea using lovastatin without any negative effect on cellulolytic bacteria [15]. In addition, naturally produced lovastatin has been shown to mitigate methane gas emissions, while simultaneously enhancing digestibility of feed [16].

A previous study has reported the effects of naturally produced lovastatin from fermented-Monascus purpureus red rice powder on cattle [17]. High dose of fermented-Monascus purpureus red rice powder (100g/day and above) supplementation adversely affected dry matter intake and ruminant physiology. We have recently reported that supplementation of naturally produced lovastatin in goats as being capable of mitigating methane emissions effectively without adversely affecting digestion and rumen fermentation, except that animals fed the highest level (6 mg/kg BW) had lower appetite [18]. Statins have been associated with the development of muscle-related adverse effects, however the effects of lovastatin on the skeletal muscle in goats have not been studied. Therefore, this follow-up study was conducted to examine the effects of lovastatin on the histology and proteome profile of the goat skeletal muscle from the above study to further elucidate whether supplementation of lovastatin affects the health and wellbeing of the goats.

## Material and Methods

### Animals and management

This study was approved by the Animal Care and Ethics Committee of the Universiti Putra Malaysia (UPM/IACUC/AUP-R0087/2015). Detail protocols of the study have been reported [18]. Briefly, twenty intact male Saanen goatsof 4–5 months old with average live weight of 26 ± 3.4 kg were used in the 12-week feeding trial. The animals were randomly assigned in equal numbers and fed a total mixed ration containing 50% rice straw, 22.8% concentrates and 27.2% of various proportions of untreated or treated PKC to achieve the target daily intake level of 0 (Control), 2 (Low), 4 (Medium) or 6 (High) mg lovastatin/kg BW. The lovastatin was produced by solid state fermentation using PKC (palm kernel cake) and Aspergillus terreus (ATCC 74135) [18].

### Slaughtering and sample collection

At the end of feeding trial, the animals were humanely slaughtered according to the standard protocol of halal (Muslim) slaughtering (MS1500:2009). Immediately after skin removal and evisceration, longissimus thoracis et lumborum muscle was collected from each animal. The muscle samples collected for histology analysis were rinsed with normal saline solution before fixing in 10% PBS buffered formalin solution. Muscle samples were snapped frozen with liquid nitrogen for proteomics analysis and kept in −80°C until further analysis.

### Histology

Muscle samples were removed from the formalin solution, dehydrated in an increasing ethanol series and routinely processed for paraffin embedding. The samples were sectioned at 5□m and stained with haematoxilin-eosin. From each muscle, five locations were sectioned and each location was mounted on a slide. and viewed with a Leica DM LB2 upright light microscope (Leica microsystems Wetzlar GmbH, Wetzlar, Germany). Three images were captured from each slide under 20 × magnification using a Leica DFC320 digital camera connected to a computer which was controlled with Leica IM50 v4.0 software (Leica microsystems Wetzlar GmbH, Wetzlar, Germany).

The muscle tissues were evaluated for evidence of necrosis, degeneration, interstitial space and vacuolization. Numerical scores were assigned based on degree of severity (0 = normal to 5 = severe) according to Gall et al. [19] (Table 1). Histological scores between every treatment groups were compared using Kruskal-Wallis test. Statistical confidence was considered as P<0.05.

**Table 1.** Scoring system used in histological analysis on *longissimus thoracis et lumborum* muscle of goats supplemented with naturally produced lovastatin.

| Score | Histopathologic injury | Percentage |
| --- | --- | --- |
| 0 | Normal | 0% |
| 1 | Minimal | <15% |
| 2 | Mild | ≤25% |
| 3 | Moderate | ≤ 40% |
| 4 | Marked | ≥50 |

### Immunoblot analysis of myofibrillar proteins

All muscle samples were pulverized into powder form with pestle and mortar using liquid nitrogen. Myofibrillar proteins were extracted according to Morzel et al. [20] with slight modifications. Approximately 100mg of muscle powder was mixed with 1ml of extraction buffer containing 150mM NaCl, 25mM KCl, 3mM MgCl2, 4mM EDTA and5μlof protease inhibitor (Calbiochem®) at pH6.5. The mixture was vortexed for 30s before centrifuging at 500rpm for 5 min. After that, the supernatant was transferred to a new tube, and centrifuged again at 4340 rpm for 15 mins at 4°C. The resulting pellet was washed twice with 1ml of 50mM KCl solution at pH6.4 and once with 1ml of 20mM phosphate buffer at pH6. The pellet was eventually suspended in Tris-SDS buffer.

The extracted myofibrillar proteins were separated by sodium dodecyl sulphate-polyacrylamide gel electrophoresis (SDS-PAGE) [21]. The proteins were mixed with sample loading buffer (62.5mM Tris-HCl at pH6.8, 25% glycerol, 2% SDS, 0.01% (w/v) bromophenol blue and 5% □-mercaptoethanol) at ratio of 1: 5 and incubated at 94°C for 5 min. For myosin heavy chain, the proteins were separated with 4% stacking and 5% resolving gels. Whilst, other proteins were separated with 4% stacking and 12% resolving gels for troponin T. The SDS-PAGE was conducted at a constant current of 15mA/gel for 15 min, followed by 20mA/gel until the bromophenol blue dye reached the bottom of the gel.

Following electrophoresis, the gel was equilibrated in the transfer buffer (25mM Tris, 192mM glycine and 20% (v/v) methanol at pH 8.3) for 10 min before blotted for 2 hr at 250mA with voltage limit of 25 V onto a polyvinylidene difluoride membrane. Membranes were blocked with 5% bovine serum albumin in TBST buffer (100mM Tris-HCl, 150mM NaCl, 0.05% Tween 20 and 0.05% SDS) overnight and incubated with primary antibody. For myosin heavy chain, the membranes were incubated with 1:500 dilution of monoclonal anti-myosin (skeletal, fast) (M4276, Sigma-Aldrich, USA) and monoclonal anti-myosin (skeletal, slow) (M8421, Sigma-Aldrich, USA) antibodies in 3% BSA in TBST buffer. Monoclonal anti-troponin-T antibody (T6277, Sigma-Aldrich, USA) was used as the primary antibody for troponin-T. After that, the membranes were washed with TBST buffer for three times and subsequently incubated in 1:10,000 dilution of secondary antibody (anti-mouse IgG whole molecule conjugated with alkaline phosphatase) (A3562, Sigma-Aldrich, USA). The membranes were then washed with TBST buffer for three times and detected using AP detection kit (Bio-Rad, USA). Protein bands were visualized using Gel DocTM XR+ System (Bio-Rad, USA) and quantified with Image LabTM software (Bio-Rad, USA).

The difference in the band intensity of myosin heavy chain and actin amongst the treatment groups were determined statistically using one-way analysis of variance (ANOVA) at 95% confidence level. The data analysis was conducted using SAS software (version 9.1, SAS Inst. Inc., U.S.A.). The data is presented as means ± standard error of means (S.E.M.).

### Liquid chromatography mass spectrometry

Crude protein was extracted from each muscle sample. Briefly, 0.2 g of muscle sample in powder form was mixed with 1ml of cold buffer containing 100mM Tris, pH 8.3 and 10 µl protease inhibitor (Calbiochem®). The samples were mixed thoroughly with vortex for 30s and centrifuged at 4°µC for 20min at 15,000 g. The supernatant was carefully collected and the concentration was determined using the Bradford assay [22].

Each protein sample (100°g) was reduced with 50mM DTT at 60°C for 60min and alkylated with 50mM iodoacetamide in the dark for 45 min at room temperature. Then, proteins were diluted with 50mM ammonium bicarbonate and digested with trypsin at 37°C overnight. The digestion process was stopped by adding 0.5% formic acid. Digested peptides were desalted using C18 ZipTip pipette tips (Millipore, Billerica, USA) according to the supplier’s instructions, and resuspended in 0.1% formic acid.

The purified digested peptides were separated with reverse phase liquid chromatography using a Dionex Ultimate 3000 RSLCnano system (Thermo Fisher Scientific) and analyzed by tandem mass spectrometry using Orbitrap Fusion mass spectrometry (Thermo Fisher Scientific). Peptide samples (2□l) were separated on the EASY-Spray Column Acclaim PepMapTM C18 100A□ (2□m particle size, 50□m id × 25cm; Thermo Fisher Scientific) by a gradient from 5% to 40% of buffer B (0.1% formic acid in acetonitrile) at 300nL/min flow over 91 min. The remaining peptides were eluted by a short gradient from 40% to 95% buffer B in 2 min.

The eluting peptides were analyzed using tandem mass spectrometry using Orbitrap Fusion mass spectrometry. Full scan spectra were collected using the following parameters: scan range 310-1800 m/z, resolving power of 120,000, AGC target of 4.0 e5 (400,000), and maximum injection time of 50 ms. The method consisted of 3 s Top Speed Mode where precursors were selected for a maximum 3 s cycle. Only precursors with an assigned monoisotopic m/z and a charge state of 2 – 7 were further analyzed for MS2. All precursors were filtered using a 20 second dynamic exclusion window and intensity threshold of 5000. The MS2 spectra were analyzes using the following parameters: rapid scan rate with a resolving power of 60,000, AGC target of 1.0e2 (100), 1.6 m/z isolation window, and a maximum injection time of 250 ms. Precursors were fragmented by CID and HCD at normalized collision energy of 30% and 28%.

The raw data obtained was analyzed using Thermo ScientificTM Proteome DiscovererTM Software Version 2.1 (Thermo Fisher Scientific) with searching database of goat (Capra hircus) and mammalian downloaded from UniProt. The parameters for searching were set as follows: missed cleavage: 2; MS1 tolerance: 10ppm; MS2 tolerance: 0.6Da; variable modification: oxidation (M), deamidation of aspargine (N) and glutamine (Q); and fixed modification: carbamidomethyl (C). All peptides were validated using the percolator© algorithm based on q-value less than 1% false discovery rate.

Quantitative analysis of the data was analyzed using Perseus version 1.6.0.2 to identify the differentially expressed proteins in the muscle between the control and lovastatin-treated groups. Pair-wise comparison between control and each lovastatin-treated group was conducted using Student T-test. Gene ontology enrichment analysis and functional annotation of the identified proteins were performed using Database for Annotation, Visualization and Integrated Discovery (DAVID) version 6.8 (https://david.ncifcrf.gov) [23]. In addition, the cell and molecular functions and canonical pathways of these proteins were identified using Ingenuity® Pathway Analysis (IPA®) (Qiagen, Germany).

## Results

### Histology

The histological examinations showed that skeletal muscle of animals supplemented with lovastatin suffered from light to moderate damage (Figure 1). Mild haemorrhage was observed in the Medium treatment group while skeletal muscle of the High treatment group showed relatively severe degeneration as compared to other treatment groups. Descriptive data for each group is shown in Table 2. The Kruskal-Wallis test showed a significant difference among the four groups. It was observed that the muscle of the Control group was normal, with no signs of any degeneration. The scores of necrosis, interstitial space and vacuolization of the Low treatment group were similar (P>0.05) to the Control group, except the score of degeneration (P<0.05). When the Medium and High treatment groups compared with the Control group, the scores of degeneration, necrosis, interstitial space and vacuolization were significantly different (P<0.05). The scores of necrosis, interstitial space and vacuolization, but not of degeneration of Low treatment group, were significantly different (P<0.05) with Medium treatment group. When comparing the Medium and High treatment groups, only the scores of necrosis and degeneration were significantly different (P<0.05) between both groups.

**Fig 1.**
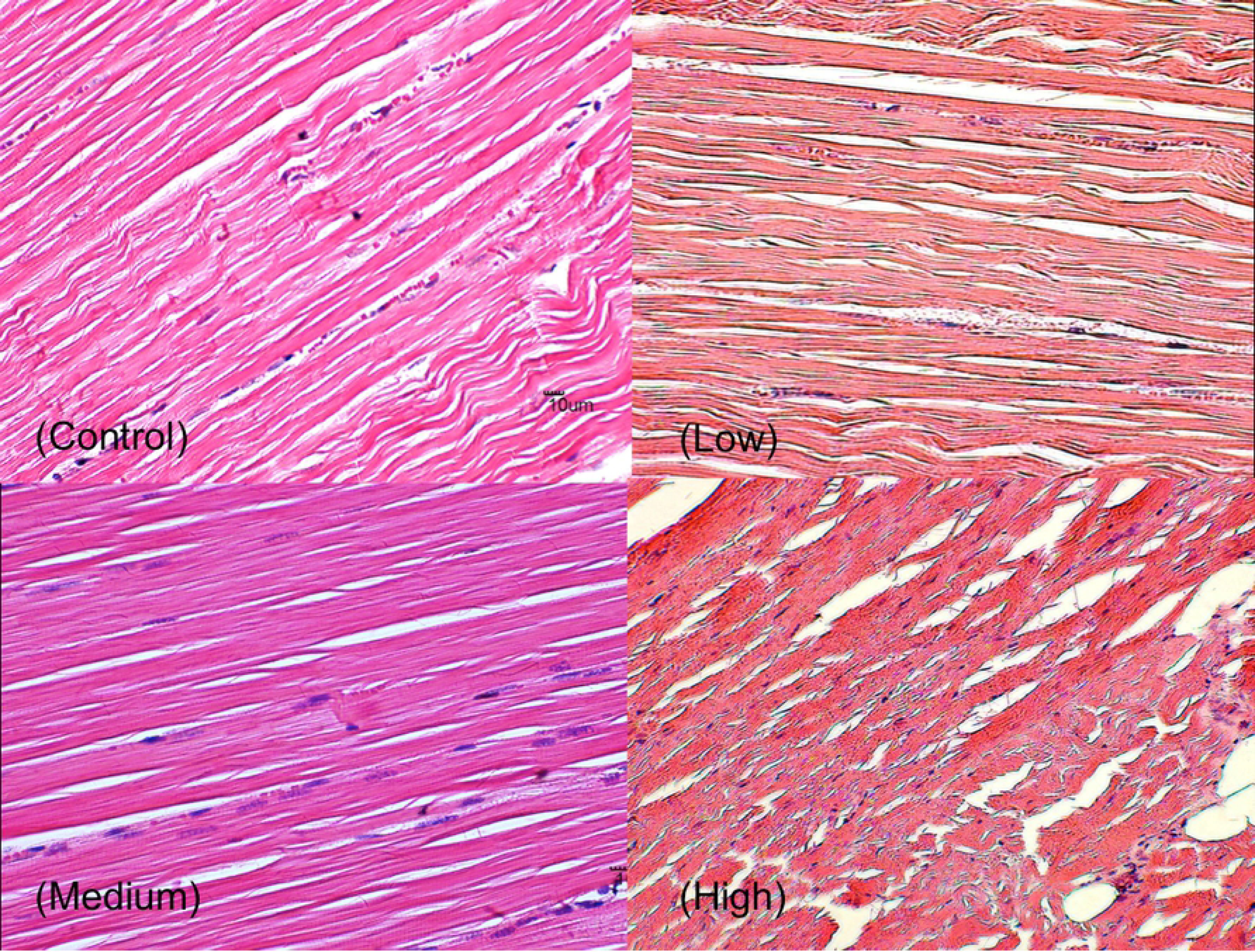
Histological analysis on longissimus thoracis et lumborum muscle of goats supplemented with naturally produced lovastatin. Control, Low, Medium and High represent 0, 2, 4 and 6 mg lovastatin/kg BW, respectively. Mild haemorrhage was observed in the Medium treatment group. Skeletal muscle of High treatment group showed relatively severe degeneration as compared to other treatment groups.

**Table 2.**
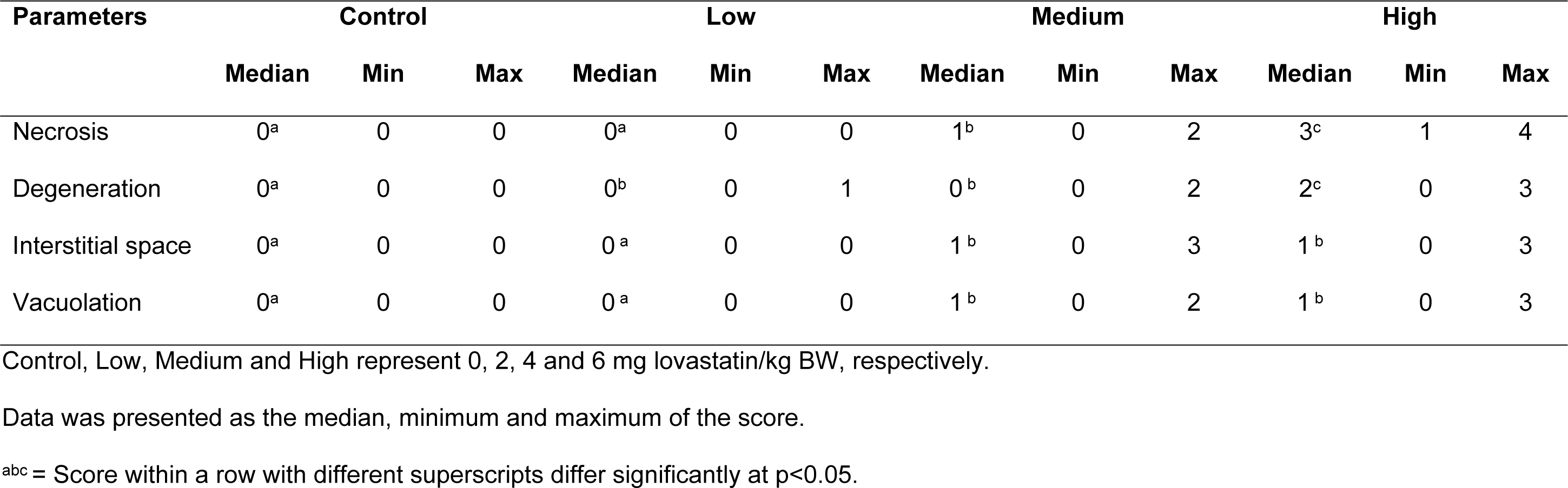
Histology scores of *longissimus thoracis et lumborum* muscle of goats supplemented with naturally produced lovastatin.

### Expressions of myosin heavy chain and actin

The difference in the immunoreactivities of myosin heavy chain and actin of longissimus thoracis et lumborum muscle in the goats supplemented with various lovastatin levels is presented in Table 3. There were significant differences in the expression of myosin heavy chain between the treatment groups, while the expression of actin was found to be unaffected by lovastatin supplementation. The immunoreactivities of myosin heavy chain in the High treatment group were significantly lower than the Control, Low and Medium treatment groups. In contrast, the immunoreactivities of myosin heavy chain were similar between the Control, Low and Medium treatment groups.

**Table 3.** Differences in immunoreactivities of myosin heavy chain and actin in *longissimus thoracis et lumborum* muscle of goats supplemented with naturally produced lovastatin.

| Protein | Control | Low | Medium | High |
| --- | --- | --- | --- | --- |
| Myosin heavy chain | 0.36±0.01<sup>a</sup> | 0.36±0.00<sup>a</sup> | 0.36±0.01<sup>a</sup> | 0.35±0.00<sup>b</sup> |
| Actin | 0.38±0.00<sup>a</sup> | 0.39±0.00<sup>a</sup> | 0.39±0.00<sup>a</sup> | 0.38±0.00<sup>a</sup> |

Control, Low, Medium and High represent 0, 2, 4 and 6 mg lovastatin/kg BW, respectively.
<sup>ab</sup> Mean within a row with different superscripts differ significantly at $p < 0.05$ .

### Expressions of myosin heavy chain and actin

The difference in the immunoreactivities of myosin heavy chain and actin of longissimus thoracis et lumborum muscle in the goats supplemented with various lovastatin levels is presented in Table 3. There were significant differences in the expression of myosin heavy chain between the treatment groups, while the expression of actin was found to be unaffected by lovastatin supplementation. The immunoreactivities of myosin heavy chain in the High treatment group were significantly lower than the Control, Low and Medium treatment groups. In contrast, the immunoreactivities of myosin heavy chain were similar between the Control, Low and Medium treatment groups.

### Differentially expressed proteins

The present study identified approximately 400 proteins in the longissimus thoracis et lumborum muscle of goat. Comparisons between control and lovastatin-treated groups demonstrated that lovastatin supplementation induced complex modifications to the protein expression patterns in the longissimus thoracis et lumborum muscle of the goat. When the Low treatment and Control groups were compared, there were 25 proteins down-regulated and five proteins up-regulated in the Low treatment group (Table 4). When the Medium treatment and Control groups were compared, 21 proteins were observed to be down-regulated and five proteins up-regulated (Table 5). When the Control and High treatment groups were compared, 23 proteins were down-regulated and one protein was up-regulated in the muscle tissue (Table 6).

**Table 4.**
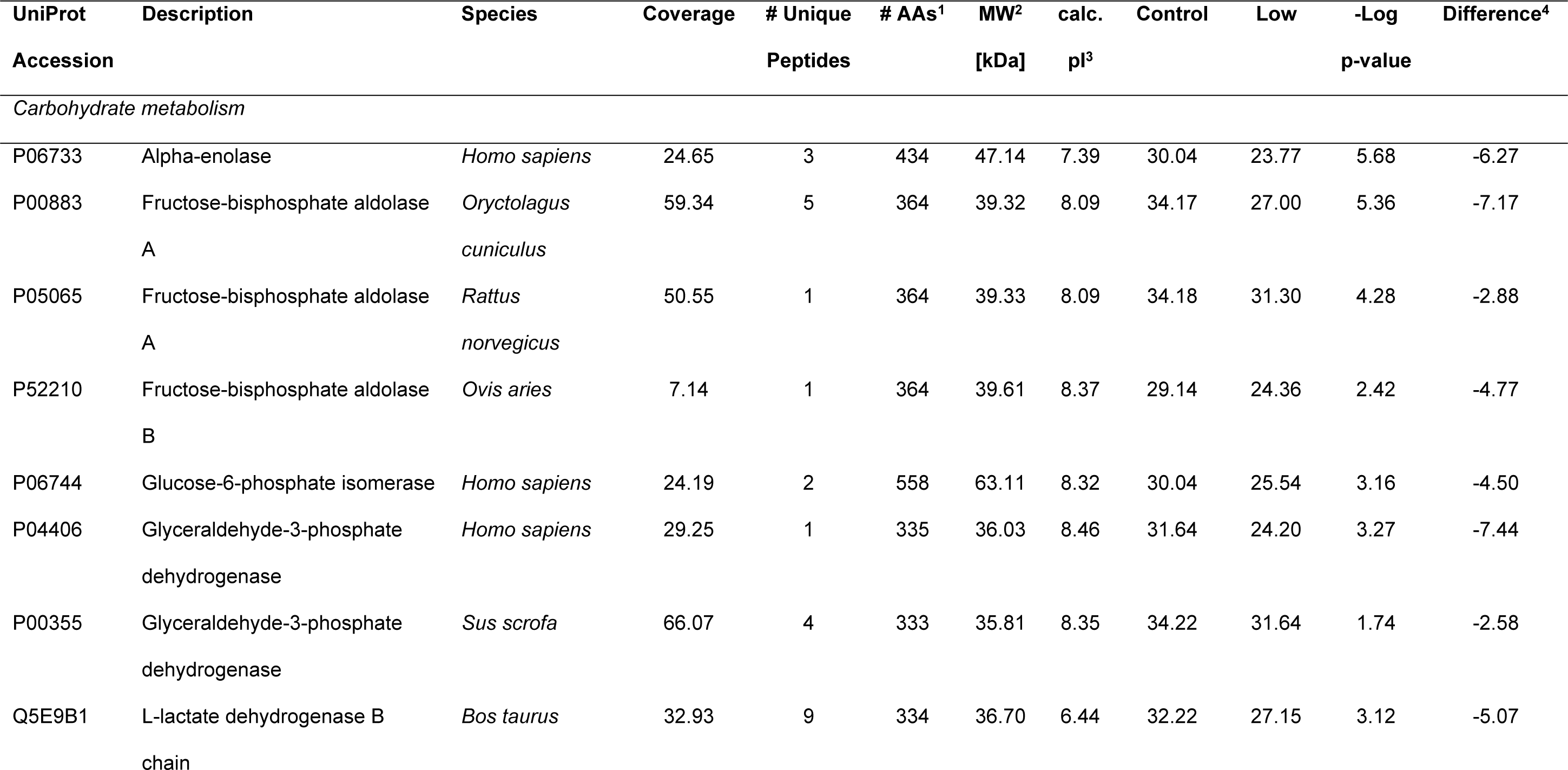
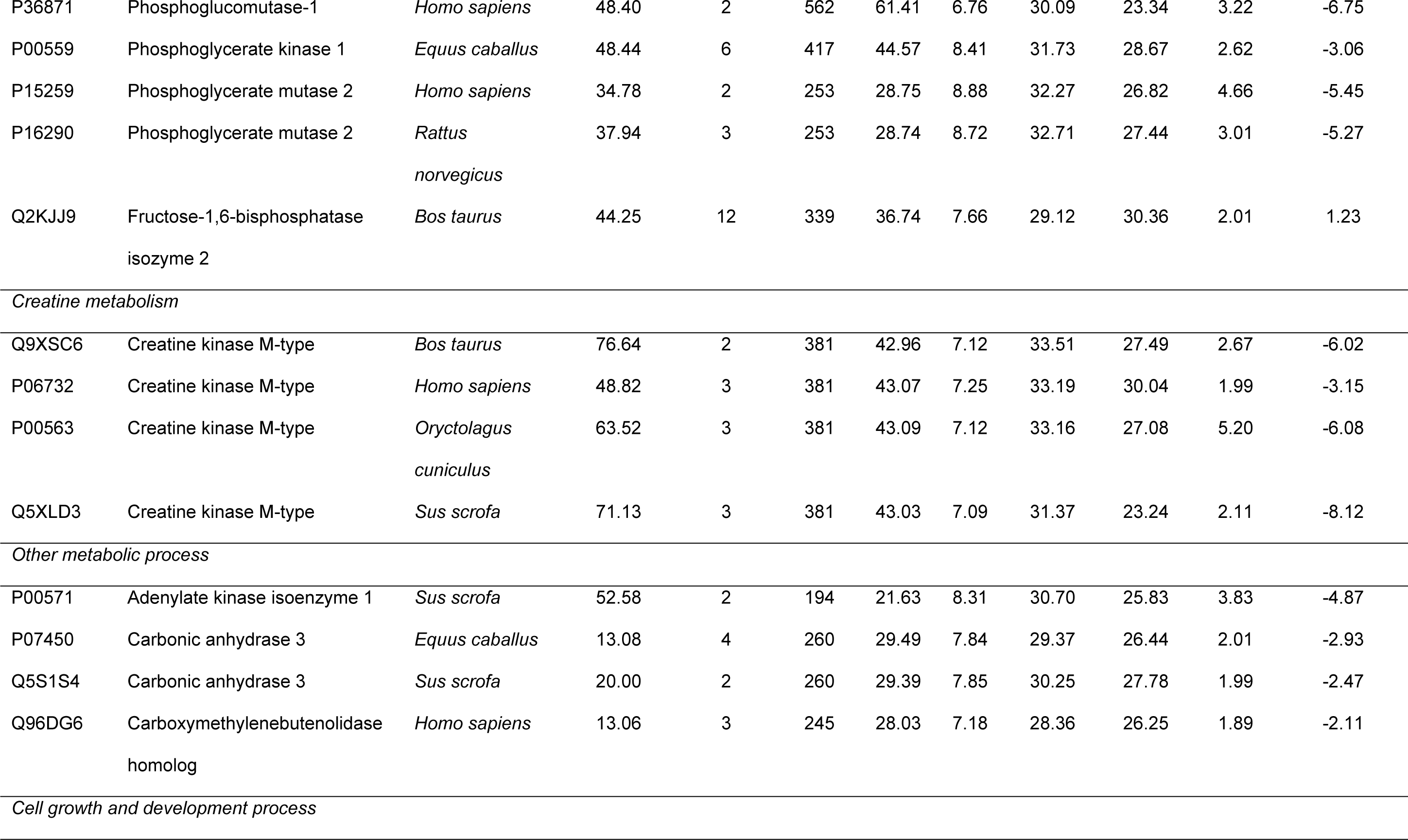
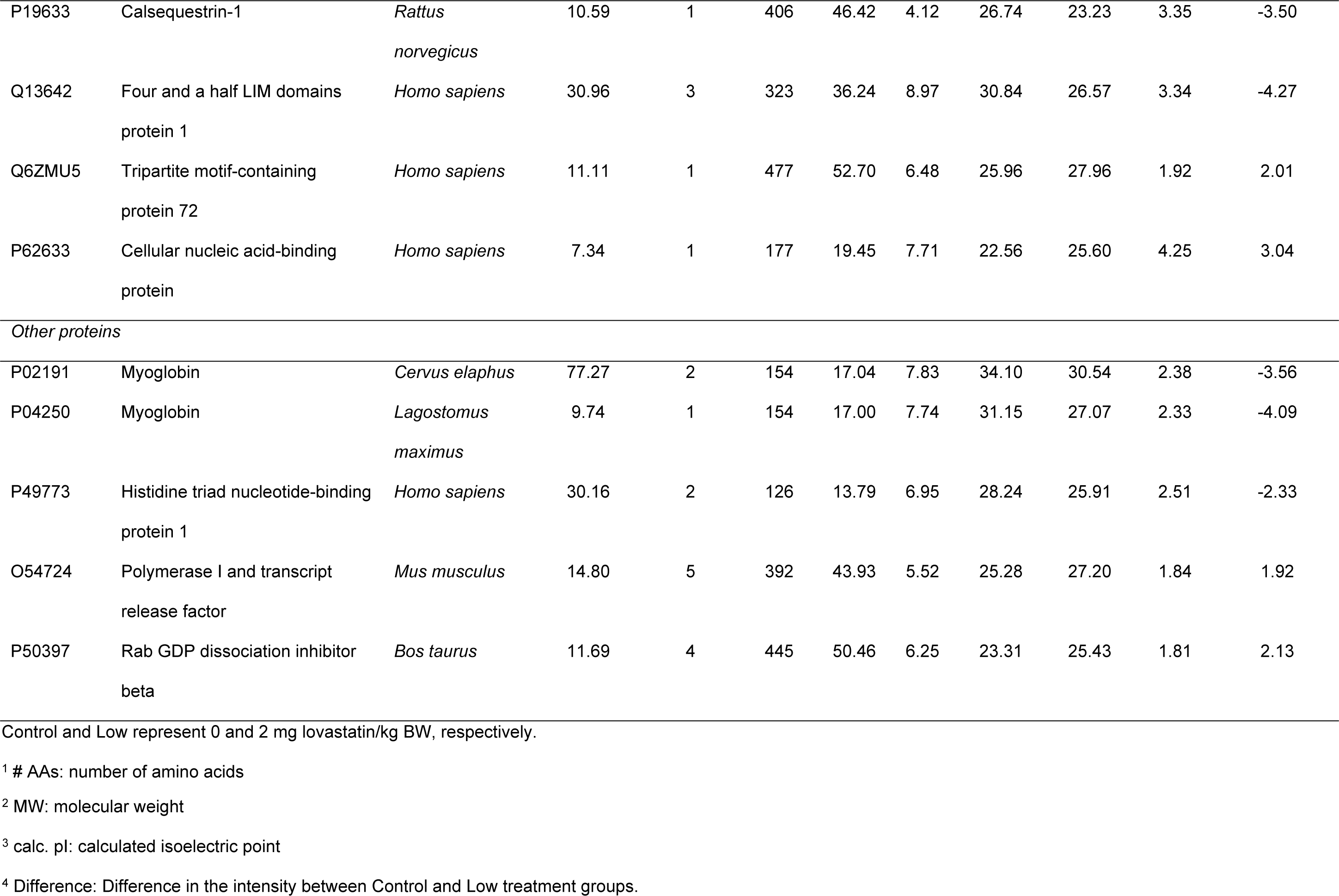
Differentially expressed proteins in *longissimus thoracis et lumborum* muscle of goats supplemented with 2mg lovastatin/ kg body weight.

**Table 5.**
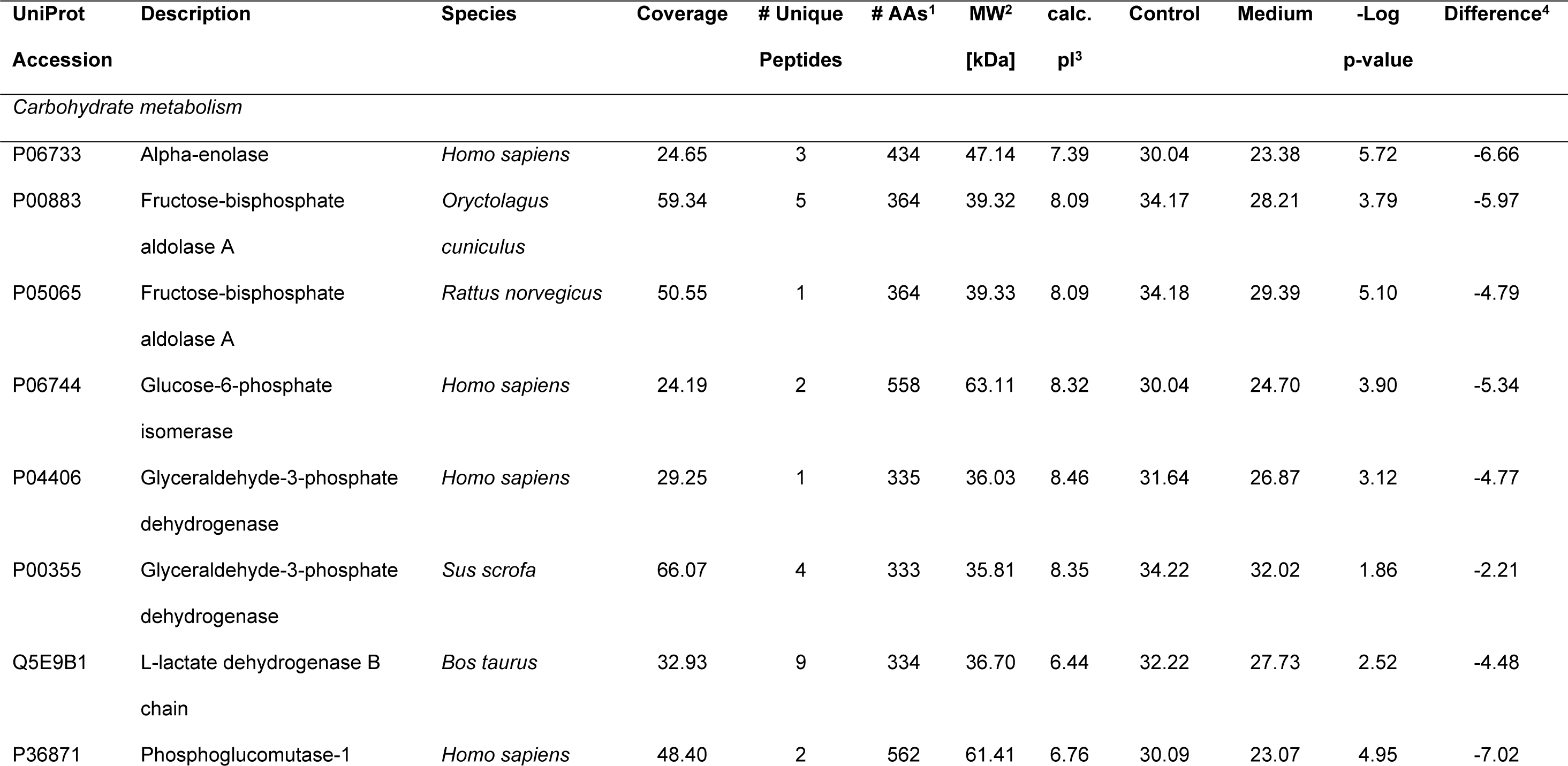
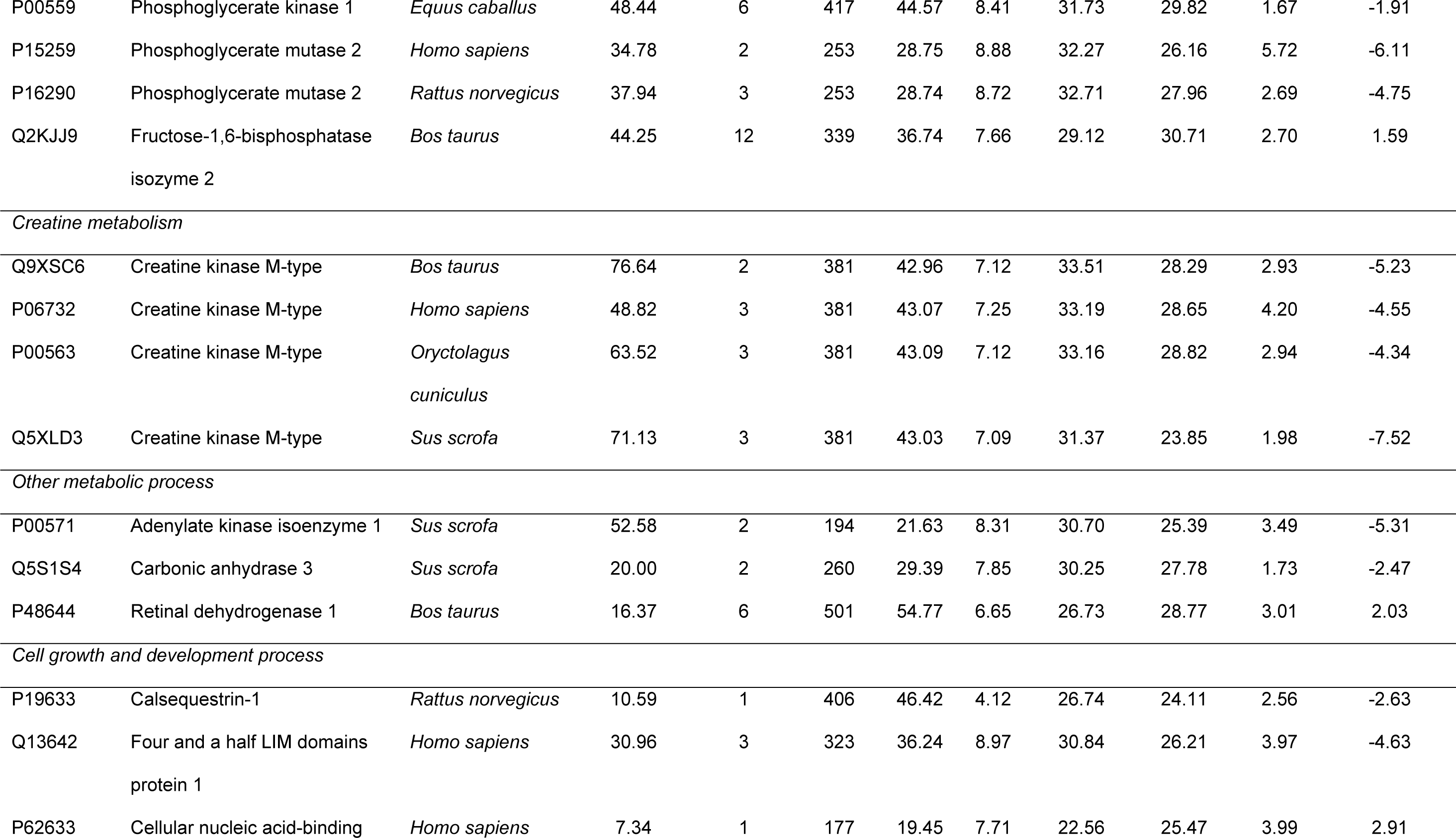
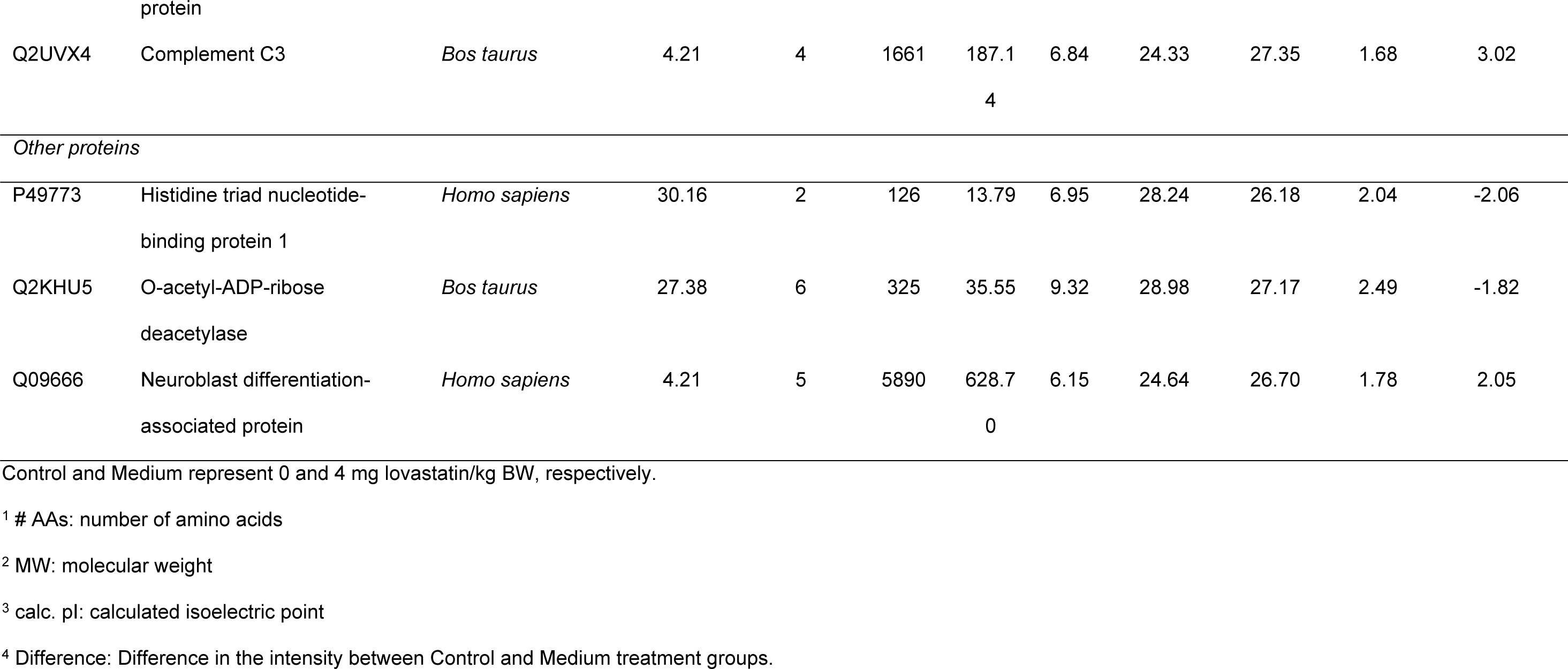
Differentially expressed proteins in *longissimus thoracis et lumborum* muscle of goats supplemented with 4mg lovastatin/ kg body weight.

**Table 6.**
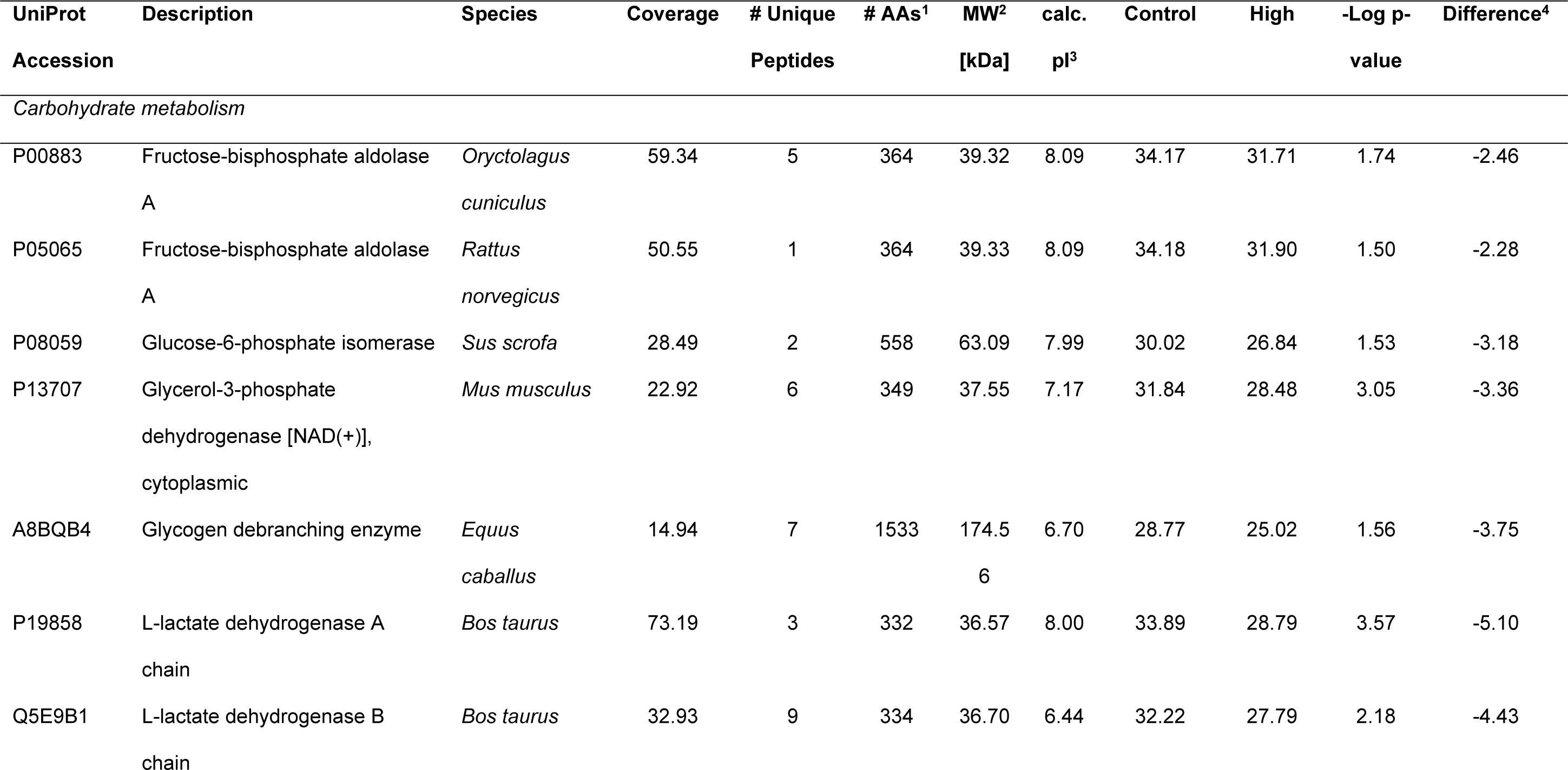
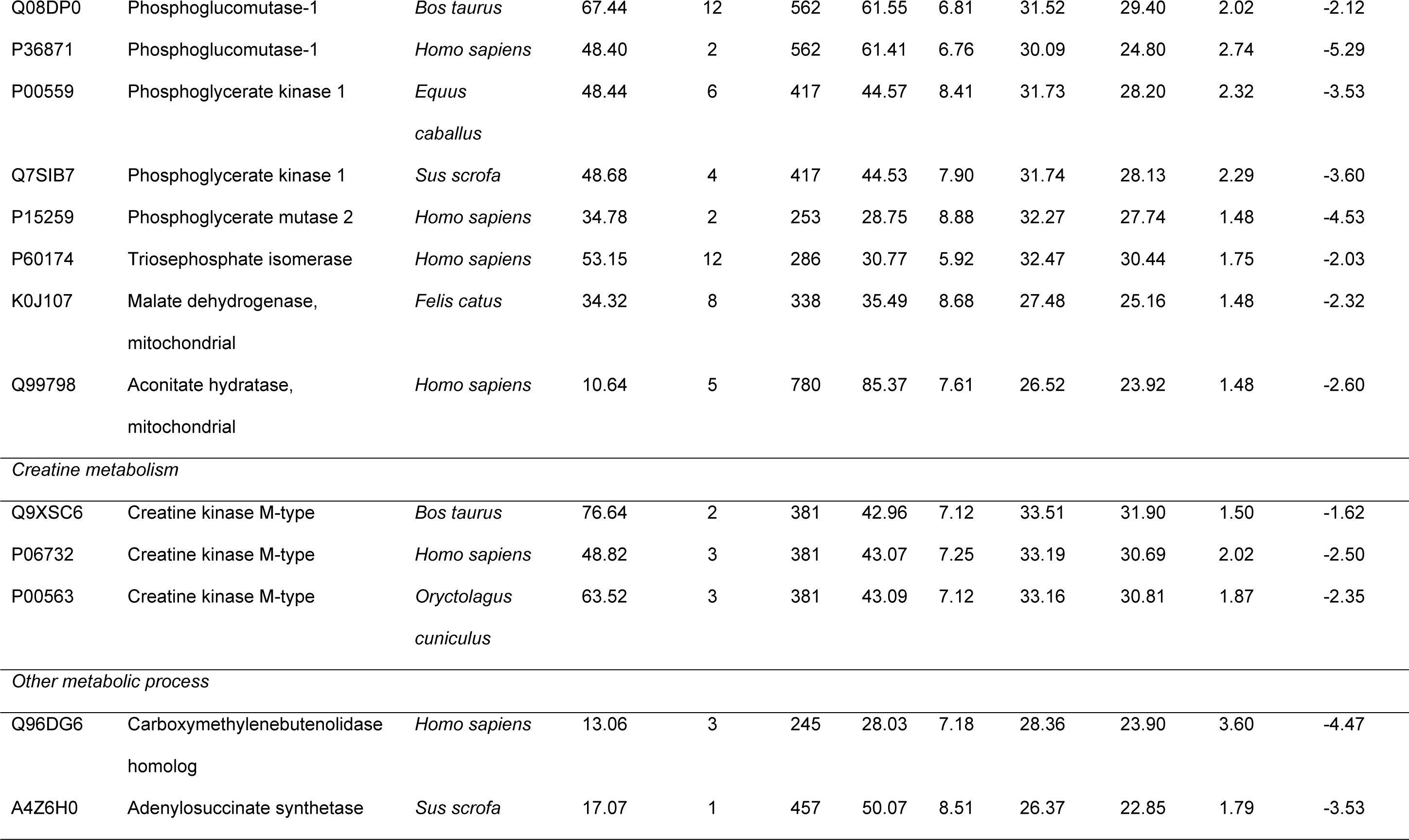
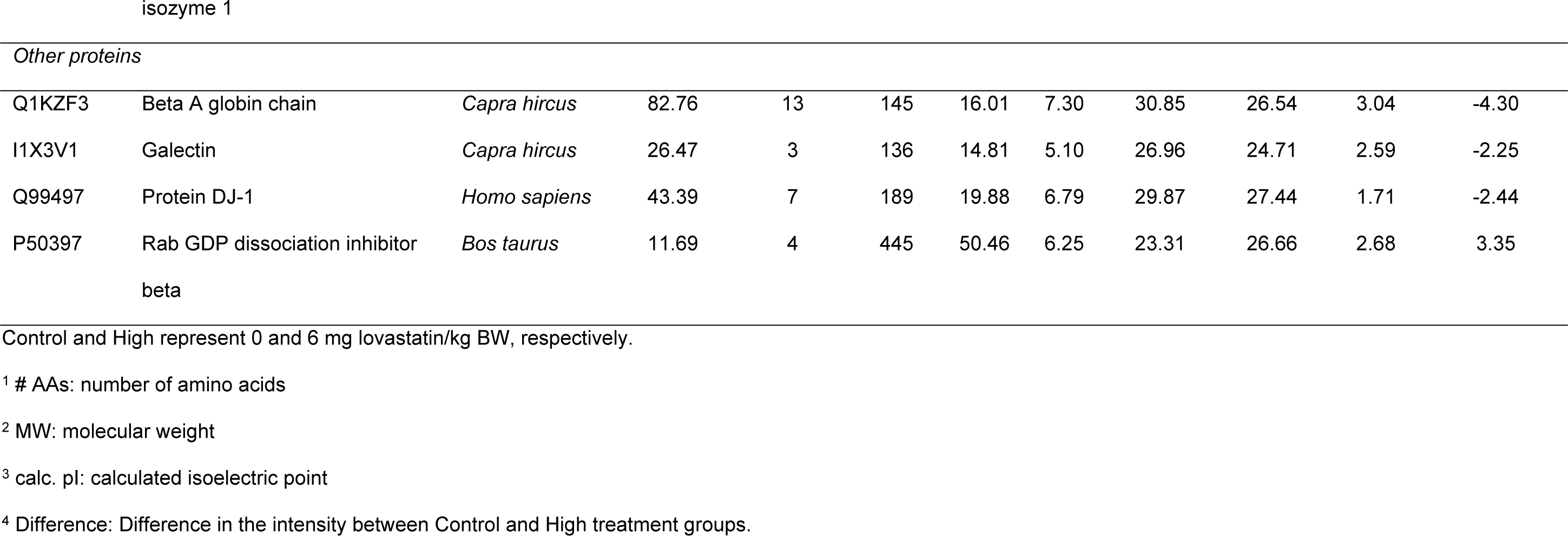
Differentially expressed proteins in *longissimus thoracis et lumborum* muscle of goats supplemented with 6mg lovastatin/ kg body weight.

The differentially expressed proteins were grouped on the basis of their functional role in the following categories: carbohydrate metabolism, creatine metabolism, other metabolic process, cell growth and development process and others. Most of the proteins affected were involved in carbohydrate metabolism, in which fructose-bisphosphate aldolase A, glucose-6-phosphate isomerase, L-lactate dehydrogenase B chain, phosphoglucomutase-1, phosphoglycerate kinase 1, and phosphoglycerate mutase 2 were down-regulated in all lovastatin treatment groups when compared to the Control group. The muscle of the Low and Medium treatment groups showed a down-regulation in the alpha-enolase and glyceraldehyde-3-phosphate dehydrogenase when compared to the Control group. Fructose-bisphosphate aldolase B was down-regulated in the Low treatment group only, while glycerol-3-phosphate dehydrogenase [NAD+], glycogen debranching enzyme, L-lactate dehydrogenase A, malate dehydrogenase, aconitate hydratase and triosephosphate isomerase were down-regulated in the High treatment group only. Simultaneously, fructose-1,6-bisphosphatase isozyme 2 was up-regulated in the Low and Medium treatment groups when compared to the Control group.

The present findings also observed the down-regulation of creatine kinase M type in all of the lovastatin treatment groups when compared to the Control group. In addition, there were a number of proteins involved in the other metabolic processes that were identified to be differentially expressed in the lovastatin treatment groups. For example, adenylate kinase isoenzyme 1 and carbonic anhydrase 3 were down-regulated in the Low and Medium treatment groups, while carboxymethylenebutenolidase homolog was found to be down-regulated in the Low and High treatment groups. Adenylosuccinate synthetase isozyme 1 was down-regulated in the High treatment group only when compared to the Control group. Retinal dehydrogenase 1 was found to be up-regulated in the Medium treatment group when compared to Control group.

There were also proteins involved in cell growth and development processes found to be differentially expressed in the present study. For instance, calsequetrin-1 and four and a half LIM domains protein 1 (FHL1) were down-regulated, while cellular nucleic acid-binding protein was up-regulated in the Low and Medium treatment groups when compared to the Control group. Tripartite motif-containing protein 72 was found to be up-regulated in the Low treatment group as compared to the Control group. Complement C3 was up-regulated in the Medium treatment group as compared to the Control group.

The expressions of a number of proteins involved in other processes were also identified to be regulated in the present study. Myoglobin and histidine triad nucleotide-binding protein 1 were down-regulated while polymerase I and transcript factor, and Rab GDP dissociation inhibitor beta were up-regulated in the Low treatment group when compared to the Control group. In addition, the Medium treatment group showed that histidine triad nucleotide-binding protein 1 and O-acetyl-ADP-ribose deacetylase were down-regulated, while neuroblast differentiation-associated protein was up-regulated. In the High treatment group, beta A globin chain, galectin and protein DJ-1 were down-regulated while Rab GDP dissociation inhibitor beta was up-regulated.

The differentially expressed proteins were subjected to IPA® analysis. These proteins were identified to be associated with several possible canonical pathways, which including glycolysis I, gluconeogenesis I, glucose and glucose-1-phosphate degradation, creatine-phosphate biosynthesis and pyruvate fermentation to lactate. Generally, these proteins were involved in a network of carbohydrate metabolism, energy production, and skeletal and muscular system development and function.

## Discussion

Statins are the most widely used lipid lowering agents which inhibit HMG-CoA reductase in the cholesterol biosynthesis pathway. Nevertheless, the use of statins is reported to have adverse effects such as muscular pain, cramps and/or stiffness on skeletal muscles in humans [24]. We had previously reported that lovastatin effectively decreased methane production in goats [18]. Given the potential for myopathies at higher than recommended therapeutic doses the effects of naturally produced statins on the skeletal muscles of ruminants was a primary interest of ours. Hence, the present study examined the effects of naturally produced lovastatin on the histology and proteome profile of the representative goat skeletal muscle longissimus thoracis et lumborum.

### Histology

Histological examination in the present study showed that supplementation of lovastatin had light to moderate adverse effects on the longissimus thoracis et lumborum muscle. The dose levels of the lovastatin supplementation are positively correlated to the extent of cellular damage on the skeletal muscle as reported previously [25]. Supplementation of 2mg lovastatin/kg BW induced the a low but noticeable degeneration of the muscle fiber in the goats. At higher dosages (4 and 6 mg/kg) naturally produced statin resulted in the higher degree of necrosis and degeneration, as well as larger interstitial space and vacuolization in the skeletal muscle.

### Expression of myosin heavy chain and actin

Statin supplementation is reported to be involved in a large augmentation of reactive oxygen species production in skeletal muscle which induce mitochondrial impairments, reduce biogenesis mechanisms and result in muscular pain or myopathy [26]. Production of reactive oxygen species in the skeletal muscle may oxidize and degrade the myofibrillar proteins. Myosin heavy chain is the protein most sensitive to oxidation among the myofibrillar proteins [27]. Previous studies observed a decrease in the concentration of myosin heavy chain following the exposure to oxidants [20, 28]. High oxidative conditions cause cross-linking, polymerization and aggregate formation in myosin heavy chain through the formation of disulfide bonds, bityrosine and carbonyl derivatives [20]. In the present study, western blot analysis showed a slight degradation of myosin heavy chain only in the High treatment group. Although histological examination found that there was cellular damage in the Medium treatment group, there was no degradation on the myosin heavy chain detected. On the other hand, actin has been reported to be relatively stable under oxidative conditions [20, 28]. This may be attributable to the inaccessibility of oxidation sites, in which myofibrillar suspensions may be masked by the interaction of actin and myosin chains [28]. The present study also demonstrated that lovastatin supplementation had no effect on the actin of skeletal muscle in goat.

### Differentially expressed proteins

The present proteomics analysis demonstrated that supplementation of lovastatin induced modifications on the expression of a number of proteins, regardless the concentration of lovastatin. This data suggests that lovastatin had an effect on a wide range of biological functions in the muscle. Lovastatin supplementation had impaired the energy production system in the skeletal muscle, particularly in the metabolism of carbohydrate and creatine. Similar observations have been reported on the extensor digitorum longus muscle of rats treated for 2-months of 10mg atorvastatin/kg BW and 20mg fluvastatin/kg BW [29]. The impairment in the energy production system could play a major role in the development of muscle damage, which is consistent with the adverse effects observed on the longissimus thoracis et lumborum muscle through histological examination.

#### Carbohydrate metabolism

The present study showed that lovastatin supplementation down-regulated proteins involving in glycolysis, gluconeogenesis and fructose metabolism. For examples, alpha-enolase, fructose-bisphosphate aldolase A, fructose-bisphosphate aldolase B, glucose-6-phosphate isomerase, glyceraldehyde-3-phosphate dehydrogenase, phosphoglycerate kinase 1, phosphoglycerate mutase 2 and triosephosphate isomerase are glycolytic enzymes which were down-regulated following lovastatin supplementation. In addition, phosphoglucomutase-1 that catalyzing the bi-directional inter-conversion of glucose-1-phosphate and glucose-6-phosphate, was also down-regulated following lovastatin supplementation. Glucose-1-phosphate is a substrate for synthesis of UDP-glucose used to synthesis a variety of cellular constituents, while glucose-6-phosphate is the first intermediate in glycolysis.

Glycolysis is an oxygen independent pathway that converts 6-carbon glucose into pyruvate. Through this metabolic process, high energy adenosine triphosphate (ATP) molecules and reduced nicotinamide adenine dinucleotide (NADH) are generated. Similar observations of down-regulation of glycolytic enzymes in the skeletal muscle have also reported previously in rat and the down-regulation of glycolytic enzymes is a symptom of energy production failure, and can contribute to muscle damage [29]. In humans, hereditary muscle glycogenoses are characterized by defective glycolytic enzymes and leads to different degree of myopathy [30].

Interestingly, glycogen debranching enzyme, mitochondrial malate dehydrogenase and mitochondrial aconitase hydratase were identified to be down-regulated in the High treatment group only when compared to the Control group. Glycogen debranching enzyme is an important regulatory enzyme in cellular glucose utilization and energy homeostasis. This bi-functional enzyme exhibiting both of oligotransferase (oligo-1,4 to 1,4-glucantransferase, EC 2.4.1.25) and glucosidase (amylo-1,6-glucosidase, EC 3.2.1.33) activities in a single polypeptide chain. Along with phosphorylase, this enzyme catalyzes the complete degradation of glycogen and the release of glucose-1-phosphate and glucose [31]. Down-regulation of this enzyme indicates the reduction of glucose degraded from glycogen, and would subsequently affect the rate of glycolysis in the skeletal muscle.

Both, mitochondrial malate dehydrogenase and aconitate hydratase are integral components in the tricarboxylic acid (TCA) cycle which playing crucial roles in energy production and biosynthesis. Malate dehydrogenase reversibly catalyzes the oxidation of malate to oxaloacetate, which is crucial in regenerating oxaloacetate that can be utilized in the TCA cycle and amino acid production [32]. Meanwhile, aconitate hydratase catalyzes the inter-conversion of citrate, isocitrate and cis-aconitate in the TCA cycle [33]. Mitochondrial aconitate hydratase is also involved in iron metabolism and is very sensitive to reactive oxygen species [34]. Large augmentation of reactive oxygen species is produced in the skeletal muscle and is associated with the induction of mitochondrial impairment [26]. The present findings indicate that a high level of lovastatin supplementation (6mg/kg BW) results in the impairment of mitochondrial function, which is in agreement with Päivä et al. [35] reporting reduction of mitochondria volume in skeletal muscle following aggressive statin treatment.

Meanwhile, fructose-1,6-bisphosphate isozyme 2 was up-regulated in the Low and Medium treatment groups. This enzyme catalyzes the hydrolysis of fructose-1,6-bisphosphate to fructose-6-phosphate in the gluconeogenesis [36]. In glycolysis, fructose-6-phosphate is converted to fructose-1,6-bisphosphate by phosphofructokinase. This step is one of the rate-limiting steps in the glycolytic pathway. Up-regulation of fructose-1,6-bisphosphate isozyme 2 and the suppression of glycolysis prevent the breaking down of glucose, and subsequently the reduction in ATP production.

#### Creatine metabolism

Generally, statin supplementation is associated with the higher concentration of creatine kinase in the blood plasma. The present study observed a down-regulation of creatine kinase following the lovastatin supplementation. Similarly, decrease of creatine kinase in the skeletal muscle of goat was also observed in rat [29]. Creatine kinase is an enzyme catalyzes the conversion of creatine to phosphocreatine by utilizing ATP. This enzyme also catalyzes the reverse reaction to produce phosphocreatine and ATP. In tissues utilizing a large amount of ATP such as skeletal muscle, creatine kinase/phosphocreatine system plays a complex and multi-faceted role in regulating cellular energy homeostasis [37]. The ATP regeneration capacity of creatine kinase is very high and considerably exceeds both cellular utilization and replenishment through glycolysis and oxidative phosphorylation [37]. Interestingly, previous study showed that the transgenic mice lacking either the cytoplasmic or mitochondrial creatine kinase may develop muscle atrophy [38]. Together with proteins involved in carbohydrate metabolism, down-regulation of creatine kinase indicated an impairment to the energy production system, which may develop statin myopathy.

#### Other metabolic process

Down-regulation of adenylate kinase isozyme 1, adenylosuccinate synthethase isozyme 1 and carbonic anhydrase 3 may impair the energy production in the skeletal muscle. Adenylate kinase isozyme 1 and adenylosuccinate synthethase isozyme 1 play an important role in cellular energy homeostasis, and more specifically in adenine nucleotide metabolism. Adenylate kinase isozyme 1 catalyzes the reversible transfer of phosphate between ATP and adenosine monophosphate (AMP), while adenylosuccinate synthethase isozyme 1 interconverts inosine monophosphate (IMP) and AMP to regulate nucleotide levels in the tissue, and which contributes to the regulation of glycolysis. Meanwhile, the lack of carbonic anhydrase 3 is suggested to impair mitochondrial ATP synthesis in the gastrocnemius muscle of rat [39]. Furthermore, carbonic anhydrase 3 is also shown to provide protection to the cells against free radicals [40].

Retinal dehydrogenase 1, glycerol-3-phosphate dehydrogenase [NAD+], L-lactate dehydrogenase A chain and L-lactate dehydrogenase B chain are involved in redox cofactor metabolism which plays a central role in meeting cellular redox requirements of proliferating mammalian cells. Retinal dehydrogenase 1 converts retinaldehyde to retinoic acid, which directly catalyzes the regeneration of NADH. Glycerol-3-phosphate dehydrogenase catalyzes the reversible conversion of dihydroxyacetone phosphate to glycerol-3-phosphate, which involves the redox reaction of NADH and NAD+. Meanwhile, lactate dehydrogenase catalyzes the conversion of pyruvate into lactate. Usually, a large amount of lactate is generated in proliferating cells to allow high glycolytic flux to support the generation of ATP and biosynthetic precursors [41]. At the same time, generation of lactate also involves the conversion of NADH to NAD+ by lactate dehydrogenase. NAD+ is crucial as it is directly used to oxidize precursors of some nucleotides and amino acids, and also many intermediates of NAD+-dependent pathway are important precursors for biosynthesis [42]. Reduction in these proteins may affect the NADP dependent pathways, then blocking the ATP production pathways.

#### Cell growth and development process

Reduction in the expression of calsequestrin and FHL1, that are involved in muscle development, was associated with the muscle damage. Calsequestrin is a Ca2+-binding protein, which has been showed to be decreased in dystrophic mouse skeletal muscle [43], while mutation in FHL1 gene is associated with myopathy [44]. FHL1 is a multifunctional protein likely to be involved in ion channel binding and muscle development. Furthermore, transport proteins such as beta A globin chain, myoglobin and polymerase I and transcript release factor were also down-regulated following lovastatin supplementation. Polymerase I and transcript release factor is a protein associated with processes of vesicular transport and cholesterol homeostasis [45].

The present findings also identified proteins involved in the muscle tissue development (such as complement C3, tripartite motif-containing proteins 72, and cellular nucleic acid binding protein) which were up-regulated following lovastatin supplementation. Complement C3 is shown to be activated in skeletal muscle injury and plays a key role in the regeneration of muscle tissue [46, 47]. Tripartite motif-containing proteins are expressed in the skeletal muscle to regulate muscle coordination, atrophy and repair. This protein plays an vital role as a negative regulator of IGF-induced muscle differentiation [48]. Meanwhile, cellular nucleic acid binding protein has been reported recently that its modifications which are indicated to play a role in myotonic dystrophy type 2 disease might result in muscle atrophy through affecting myofiber membrane function [49]. Together with Rab GDP dissociation inhibitor beta which involved in the regulation of vesicle-mediated cellular transport [50], up-regulation of these proteins in the present study indicated that the mechanism associated with tissue regeneration or repair was activated in the muscle tissue.

#### Other proteins

In addition to the energy production system, proteins involved in other cellular processes were also affected by lovastatin. Proteins such as galectins, histidine triad nucleotide-binding protein 1 and Protein DJ-1, which are in involved in the regulation of the apoptotic pathway, were also down-regulated following lovastatin supplementation. Galectins have a diverse range of biological functions including regulation of pre-mRNA splicing, cell adhesion, cell growth, differentiation, apoptosis and cell cycle [51], while protein DJ-1 plays an important role in cell protection against oxidative stress and cell death [52].

Taken together, the present study has shown that lovastatin supplementation down-regulates proteins involved in the energy production system (particularly the glycolytic pathway and creatine metabolism), regardless of the concentration of lovastatin. Supplementation with a high concentration of lovastatin (6mg/ kg BW) could further impair the TCA cycle. Moreover, supplementation of lovastatin also activates tissue regeneration or repair in the muscle tissue. Furthermore, changes in the expression of proteins involved in apoptosis and oxidative damage suggests an accentuated sensitivity of statin-treated muscle to oxidative stress. Oxidative stress can promote increased proteolysis and depress protein synthesis, and trigger many conditions associated with muscle wasting [53]. Such perturbation in energy metabolism and ATP synthesis may have profound effects on protein synthesis and contribute to metabolic stress, which could play a major role in the development of myopathy.

## Conclusions

Histology scores indicated increasing muscle damage to the longissimus thoracis et lumborum muscle of goats supplemented with increasing dosages, particularly at 6mg/kg BW, of naturally produced lovastatin. In addition, western blot analysis indicated that the immunoreactivity of myosin heavy chain was only degraded in the muscle of goats supplemented with 6mg lovastatin/kg BW. Proteomics analysis revealed that lovastatin supplementation induced complex modifications to the carbohydrate metabolism, energy production, and skeletal and muscular system development of skeletal muscle of goats which may have contributed to the observed skeletal muscle damage. Putting together results of the above three analyses, it is clear that supplementation of naturally-produced lovastatin at 6 mg/kg BW is too high, which can adversely affecting health and wellbeing of the animals.

## Acknowledgements

This study was funded by the New Zealand Government to support the objectives of Livestock Research Group of the Global Research Alliance on Agricultural Greenhouse Gases and Universiti Putra Malaysia. The authors acknowledged the contribution of Proteomics Core Facility, Malaysia Genome Institute, National Institutes of Biotechnology Malaysia (NIBM).

